# Decoding Motivational States and Craving through Electrical Markers for Neural ‘Mind Reading’

**DOI:** 10.1101/2024.11.22.624825

**Authors:** Alice Mado Proverbio, Alice Zanetti

**Affiliations:** Cognitive Electrophysiology lab, Dept. of Psychology, University of Milano-Bicocca

**Keywords:** ERPs, Imagery, BCI, mental representation, mind reading, neuroimaging, LORETA, physiological state, craves

## Abstract

The aim of this electroencephalogram (EEG) study was to identify electrical neuro-markers of 12 different motivational and physiological states such as visceral craves, affective and somatosensory states, and secondary needs. Event-related potentials (ERPs) were recorded in 30 right-handed participants while recalling a specific state upon the presentation of an auditory verbal command incorporating an evocative sound background consistent with that state (e.g. the chirping of cicadas associated with the verbal complaint about feeling hot). ERP data showed larger amplitude N400 responses in the affective and somatosensory states, while the P400 component displayed greater amplitudes for the secondary and visceral states. Furthermore, the two components were also discernibly responsive to the 12 micro-categories (e.g., joy vs. pain or hunger), by providing a distinctive electric pattern for mostly all microstates. The reconstruction of the intracranial generators of surface signals revealed common imagery-related activations, including the middle and superior frontal gyri, the fusiform and lingual gyri, supramarginal, and middle occipital regions, as well as the middle temporal region. Additionally, specific regions were identified that were active for distinct mentally represented content, such as that visceral needs were associated with activations in the medial and inferior frontal gyri, uncus, precuneus, and cingulate gyrus. Affective states were associated with activations in the medial frontal, superior temporal, and middle temporal gyri. Somatosensory states (e.g., pain or cold) activated regions in the parietal cortex and the crave for music was linked to activations in the auditory and motor regions. These findings support the use of ERP markers for BCI applications.

## Introduction

In this study, we recorded brain signals associated with motivational states, needs, and cravings activated through imaginative recall to examine how mental imagery might be “recorded” through electric signals (Reddy et al., 2010). This approach offers intriguing possibilities for future advancements in technologies like “mind reading,” where brain signals could be decoded to interpret internal mental states (Leoni et al., 2024; Pereira et al., 2018), with a non-invasive technique.

Research on the neural foundations of mental imagery reveals significant overlap with perceptual processing, indicating that imaginative and perceptual experiences share a number of cognitive resources and brain regions. Central to these findings, Dijkstra et al. (2021) have shown that internally generated images share some of the basics with actual perception, although being more attenuated. Expanding on this concept, Dijkstra et al. (2022) argued that distinguishing perceptually driven experiences from self-generated imagery remains inherently challenging, as both forms of experience activate similar neural networks and can converge at later processing stages. This convergence may suggest that mental imagery functions to emulate sensory processes, lending support to the mental imagery emulation theory. This theory posits that imagination not only simulates the content of perceived stimuli but also mimics the perceptual processes themselves, aligning imagination closely with perception. From a neuroanatomical perspective, Spagna et al. (2021) proposed that frontotemporal networks, which are essential for working memory and attentional control, underpin imaginative activity. This model finds support in Dijkstra et al. (2017), who identified a top-down pathway for imagery from the inferior frontal gyrus (IFG) to the occipital cortex. Their fMRI research showed that during imagination, only this top-down pathway remains active, with heightened activation relative to perceptual processing. This elevated activation could reflect the increased attentional demands required for generating mental imagery.

Temporal dynamics further differentiate perception from imagination while highlighting shared processes. In a 2018 study, Dijkstra et al. used magnetoencephalography (MEG) to track neural activation over time, discovering that imagery does not exhibit the clear, sequential processing phases typically seen in perception. Instead, imagination is marked by more generalized, slower activation. However, overlaps in neural responses were found at specific time points especially 300 ms after stimulus onset. This pattern indicates that while initial stages of processing diverge, later stages of perception and imagination begin to converge, suggesting shared underlying functions.

Research on the neural underpinnings of mental imagery, particularly with respect to emotional and motor components, has garnered increasing scholarly attention. ERP studies have identified significant neural responses associated with the imagination of emotionally charged stimuli, revealing heightened late positive potential (LPP) responses over central areas for negative content compared to neutral or positive scenarios evoked by auditory narratives (MacNamara et al., 2018). This suggests that imagery involving emotionally intense stimulation demands more substantial neural engagement relative to neutral or positive imagery. Additional ERP research highlights further components linked to mental imagery. Ogawa and Nittono (2019), for instance, identified N400 and N700 components in anterior and centro-parietal regions during imaginability tasks, where participants were instructed to evoke mental representations in response to visually presented words. EEG and ERP methodologies have proven especially instrumental in studying motor imagery, with notable implications for medical applications within brain-computer interface (BCI) research. For example, Syrov et al. (2022) observed that during motor imagery tasks, the N200 component effectively differentiates between mental counting and motor imagery conditions, while the slow potential (SP) exhibits both longer latency and greater amplitude specifically during motor imagery, particularly in central regions (Cz electrode). Moreover, research by Mencel et al. (2022) illuminates neural patterns associated with distinct types of motor imagery, such as reaching versus grasping, with significant activation observed in frontal and centro-parietal areas, aligning with findings from Syrov et al. (2022). Collectively, these studies underscore that central areas, alongside premotor and sensorimotor cortices, are integral to the mental simulation of motor actions. The emotional dimension of mental imagery engages neural networks similar to those activated by real experiences of the same stimuli. Lang’s bio-informational theory posits that imagining an emotionally charged stimulus triggers an associative network overlapping with that involved in direct perception. Ji et al. (2016) demonstrate that imagery can evoke emotional and physiological responses (such as heart rate changes), enabling individuals to anticipate the emotional consequences of imagined events. Studies like Costa et al. (2019) reveal that imagining positive scenarios activates the medial prefrontal cortex (mPFC), nucleus accumbens, and amygdala, while negative scenarios reduce activity in the former areas while increasing amygdala activation, indicating a distinct role of these regions in processing positive imagined content.

The neural and electrical markers of imagination related to various states, such as sleep, hunger, craving, and emotional experiences, reveal important insights into brain activity. However, evidence regarding the neural markers of sleep remains inconsistent. Sleep deprivation has been associated with reduced amplitudes of N1 and P300 components in sensory and frontal cortices (Boonstra et al., 2007), suggesting diminished capacity to detect and anticipate stimuli (Trošt Bobić et al., 2016).

Regarding basic needs like hunger and thirst, specific neural markers have been identified. The need for hydration activates the anterior cingulate cortex and the insula (Adams et al., 2021; McKinley et al., 2019). Neural responses to food-related stimuli include larger amplitudes of P1, P2, P300, LPP, and N200 components, particularly when participants are hungry or when presented with more calorically dense foods (Carbine et al., 2018). Similar patterns are observed in craving responses, where addiction-related cravings for substances like cocaine or pathological gambling activate reward networks, including the prefrontal cortex, cingulate cortex, and fusiform gyrus (Antons et al., 2023). In individuals with addiction, ERP components such as N170, P2, P300 and reflect the difficulty in inhibiting cravings (Habelt et al., 2020).

Motor desire, studied through EEG techniques like movement-related cortical potentials, shows deflections in amplitude 400 ms before movement execution, with involvement of frontal, parietal, and motor-related areas (Pereira et al., 2017). EEG studies also indicate greater theta band activity during motor imagery, particularly in the SMA and dorsolateral PFC (Van der Lubbe et al., 2021). Neuroimaging studies further support these findings, revealing activity in premotor, parietal, and cerebellar regions during both motor imagery and actual movement (Hardwick et al., 2018).

The neural correlates of desires for activities like video gaming and music listening also align with patterns of brain activation observed in more fundamental survival needs. For example, craving for video games in individuals with internet gaming disorder results in increased delta, theta, and beta activity in central and parieto-occipital regions (Park et al., 2023), while healthy participants exhibit activity in the temporal cortices during social gaming desire (Della Vedova and Proverbio, 2024). Music listening, a cross-cultural and ancient human activity, activates reward circuits, including the nucleus accumbens and orbitofrontal cortex, and elicits dopamine release during anticipation (Salimpoor et al., 2011; Reybrouck & Van Dyck, 2024), besides auditory areas. Emotional experiences, including joy, sadness, and fear, also involve distinct neural signatures. Positive emotions, such as joy, are associated with enhanced activation in occipital and orbitofrontal cortices (Proverbio and Cesati, 2024), while negative emotions like fear modulate early ERP components, such as N1 and P2, in centro-parietal regions (Sperl et al., 2021). The perception of sadness triggers distinct neural responses, including a larger N2 and prolonged LPP, particularly over occipital regions (Revers et al., 2022). Sensory pain perception also evokes increased N2 and P2 components in fronto-central and temporal areas (Meng et al., 2013).

In summary, although neural markers vary across different states, there is substantial evidence that brain regions involved in motivation, sensory processing, and emotional regulation reflect the underlying mental processes related to sleep, hunger, craving, and emotions.

ERP components can serve as neural markers, enabling patients to communicate with the external world by identifying specific electrical and neural activity patterns. Since their discovery, ERPs have proven to be reliable markers for category-specific visual and auditory processing (Dijkstra et al., 2020; Proverbio et al., 2022). Several studies have used EEG to identify ERPs relevant to Brain-Computer Interface (BCI) technologies. For instance, Proverbio et al. (2023) explored neural markers through ERPs in a task involving visual and auditory imagery of concrete entities. The following peaks were identified at specific scalp sites and latencies, during imagination of infants (centroparietal positivity, CPP, and late CPP), human faces (anterior negativity, AN), animals (anterior positivity, AP), music (P300-like), speech (N400-like), affective vocalizations (P2-like) and sensory (visual vs auditory) modality (PN300). Overall, perception and imagery conditions shared some common electro/cortical markers, but during imagery the category-dependent modulation of ERPs was long latency and more anterior, with respect to the perceptual condition.

Similarly, Proverbio et al. (2023) investigated ERP components associated with imagined motivational states prompted by previously validated pictograms, aiming to provide relevant data for BCI technologies in assessing consciousness in patients with disorders and offering them communication methods. Participants recalled motivational states associated to: primary needs (hunger, thirst, sleep), affective states (happiness, sadness, fear), secondary needs (play, movement, music), and somatosensory states (pain, heat, cold). Results showed larger N400 peaks during perception than imagination across most categories, except for somatosensory states, where mental imagery elicited a stronger response than primary needs and affective states, particularly in anterior frontal sites. The Late Positive Potential (LPP) did not differ significantly between perception and imagination conditions (except for somatosensory states) and was larger over the right hemisphere. Significant differences were observed in the anterior N400 component across specific micro-categories, such as between cold and heat, happiness and fear, or play and movement. The findings derived from these studies might be of considerable importance, as advancements in signal processing may contribute to the development of “mind reading” technologies within Brain-Computer Interface (BCI) applications. In fact, algorithms have been recently developed to automatically classify these signals using deep learning approaches applied to EEG traces and ERP epochs (Leoni et al., 2021; Leoni et al., 2022; Leoni et al., 2023).

In order to determine whether the modulation of these ERP components was indeed related to the distinct nature of motivational states (and their associated neural signatures), rather than to the contingent, pictorial, content of the cues, brief auditory commands (with expressive voice and evocative background) were employed as cues in the present study. The aim was to identify universal electric markers, generalizable from the semantic point of view. Building upon previous findings from the pictogram study by Proverbio and Pischedda (2023), we hypothesized that both macro and micro-motivational states would modulate late-latency ERP components, such as the P/N400, in a distinct manner. Consistent with the predictions of Dijkstra (2017, 2021), we also anticipated that imagery would predominantly activate anterior brain regions at latencies greater than 300 ms, reflecting higher-order cognitive processes linked to mental representations.

## Methods and Procedure

### Participants

Thirty participants (17 females and 13 males) aged between 18-28 years, with no current or history of psychiatric or neurological disorders, took part in the study. Seven participants were subsequently excluded due to excessive EEG artifacts. The final sample included 23 participants, comprising 16 females and 7 males, with a mean age of 22.52 years (SE = 2.13) and an average education level of 16.3 years (SE = 1.72). All participants provided informed written consent and were right-handed according to the Edinburg inventory. They were recruited through the online SONA System platform and received ECTS credits for their participation.

The research project, entitled “Auditory imagery in BCI mental reconstruction” was pre-approved by the Research Assessment Committee of the Department of Psychology (CRIP) for minimal risk projects, under the aegis of the Ethical committee of University of Milano-Bicocca, on February 9th, 2024, protocol n: RM-2024-775).

### Stimulus and materials

Auditory stimuli were designed by combining a human expressive voice with a background sound to evoke a context related to the targeted needs. The stimuli included: primary or visceral needs (hunger, thirst, and sleep), homeostatic or somatosensory sensations (cold, heat, and pain), emotional or affective states (sadness, joy, and fear), and secondary needs (desire for music, movement, and play). 17 audio clips were recorded for each microcategory, each replicated twice: once with a male voice and once with a female voice, totaling 408 stimuli. *Audacity* software was used to combining the vocal track with a background context coherent with the verbal content. Human voices were recorded using Microphone K38 by *Hompower* (SNR = 80 dB). The semantic content, the prosodic intonation and the emotional tone of all voices were coherent and appropriately matched. Some of the background sounds were recorded using the same microphone, while others were sourced from the publicly accessible BBC Sound Effects library for scientific purposes (https://sound-effects.bbcrewind.co.uk/search).

Stimuli were balanced for both length (number of words in the spoken message) and duration (in seconds) across the classes. Length measures were analyzed using a one-way ANOVA across the 12 micro-categories, which yielded a non-significant result (F11,176 = 0.192, p = 0.55), indicating that sentence lengths did not differ across categories. Stimuli intensity was balanced, with audio volume standardized to 89 dB using *MP3Gain Express*.

### Stimulus validation

The stimuli were validated using an independent sample of 40 participants (20 males, 20 females; M = 26.15 years, SE = 4.96), drawn from the same statistical cohort as the EEG study participants. The 408 audio recordings were distributed across two validation questionnaires, each tailored to the gender of the narrating voice. Each questionnaire comprised 204 acoustic stimuli, with 17 items per subcategory. The purpose of these questionnaires was to assess the efficacy and plausibility of the audio clips in conveying their respective motivational states. The surveys were conducted using the Qualtrics online platform and disseminated via invitation links shared in WhatsApp groups at the University of Milano-Bicocca. Participants were instructed to use headphones and were tasked with listening to each audio clip, identifying the described motivational state, and evaluating the plausibility of the content and the extent to which the voice effectively expressed the associated need. Responses were recorded on a 3-point Likert scale ranging from “not very plausible” to “very plausible.” Each questionnaire required approximately 25 minutes to complete.

The questionnaire analyses demonstrated a remarkably high accuracy in identifying the content of the audio clips, with an overall mean rate of 96.33% (SE = 1.3) for correct responses. Notably, the highest accuracy rates were observed for audio stimuli representing the subcategories of thirst (98%), sleep (98%), cold (98%), happiness (98%), and fear (97%). Conversely, slightly lower accuracy rates were recorded for subcategories such as play (96%), music (96%), sadness (96%), pain (96%), hunger (95%), heat (95%), and movement (94%). Based on the validation data, only audio stimuli correctly identified in at least 70% of responses were included in the final stimulus set. Consequently, seven audio clips were excluded: four featuring a female voice and three a male voice. These excluded stimuli belonged to the subcategories of thirst (1 audio), heat (3 audios), sadness (2 audios), and movement (1 audio). On average, participants rated the acoustic stimuli as “very plausible” in 47.58% (SE = 7.13) of responses, “somewhat plausible” in 33.75% (SE = 3.33), and “not very plausible” in 18.67% (SE = 4.6). A statistical analysis was conducted to examine variance based on two factors: “motivational states” (the four macro-categories) and “plausibility” (with levels: “not very plausible,” “somewhat plausible,” and “very plausible”). The analysis revealed a significant main effect of “plausibility” (F(2, 808) = 348.75, p < .001) and a significant interaction between the two variables (F(6, 808) = 9.75, p < .001). The distribution of responses across the three plausibility levels differed significantly. Specifically, “not very plausible” was selected significantly less often than both “somewhat plausible” and “very plausible” (both p < .001). Furthermore, “very plausible” ratings were assigned significantly more frequently than “somewhat plausible” (p < .001). Post-hoc comparisons for the interaction revealed that somatosensory states were rated as “very plausible” significantly more often (p < .001) and as “not very plausible” significantly less often (p < .05) than the other macro-categories. These findings suggest that “somatosensory” auditory cues—representing states such as pain, heat, and cold—were perceived as the most realistic overall.

A repeated-measures ANOVA was conducted to examine significant differences between micro-categories and perceived plausibility. The factors analyzed included motivational states (12 micro-categories) and the “plausibility” variable, categorized into three levels: “unlikely,” “somewhat likely,” and “very likely.” The analysis revealed a significant main effect for the plausibility (F2, 792 = 3.69, p < 0.001) and a significant interaction between motivational states and plausibility levels (F22, 792 = 5.71, p < 0.001). Post-hoc analyses identified significant differences between specific micro-categories and plausibility levels (p < 0.01). The “hunger” micro-category was perceived as the least plausible, while the “cold” motivational state received the highest mean number of “very likely” responses and the lowest mean number of “unlikely” responses. Among affective states, “joy” elicited fewer “very likely” judgments, as did the secondary category of “movement desire.”

### Experimental procedure

To ensure at least 50 EEG trials per micro-category, a subset of stimuli—specifically those with fewer classification errors and higher perceived realism—were repeated, resulting in 600 stimuli across 12 categories. These audio stimuli were organized into 18 sequences, each containing 33 or 34 stimuli. Due to the rapid presentation rate, sequences were structured in consecutive blocks of 4 or 5 items from the same micro-category to aid participant recall and imagery. Care was taken to avoid repeating any micro-category more than five times and to ensure a balanced representation of male and female voices. The order of micro-categories varied across sequences, and each sequence concluded with an attention-check question referencing to previously heard stimuli. Participants began by reading and signing an informed consent form and completing a laterality questionnaire. Subsequently, a 128-electrode EEG headset was applied, with the preparation process requiring approximately 25 minutes. During this time, participants reviewed the experimental instructions and completed a training session on imagery (Table 1). The training involved closing their eyes for 10 seconds to imagine an emotional state or need—movement, thirst, warmth, or sadness—corresponding to one of the four macro-categories. Afterward, participants reported the strategies they employed to imagine the state and indicated whether they recalled past experiences or generated new mental content during the exercise.

**Table 1.**
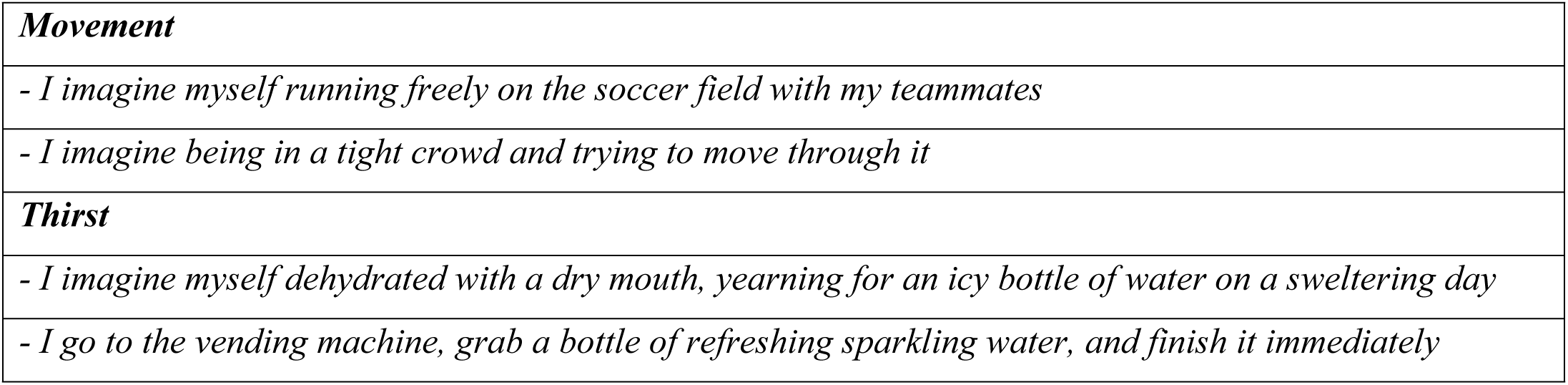
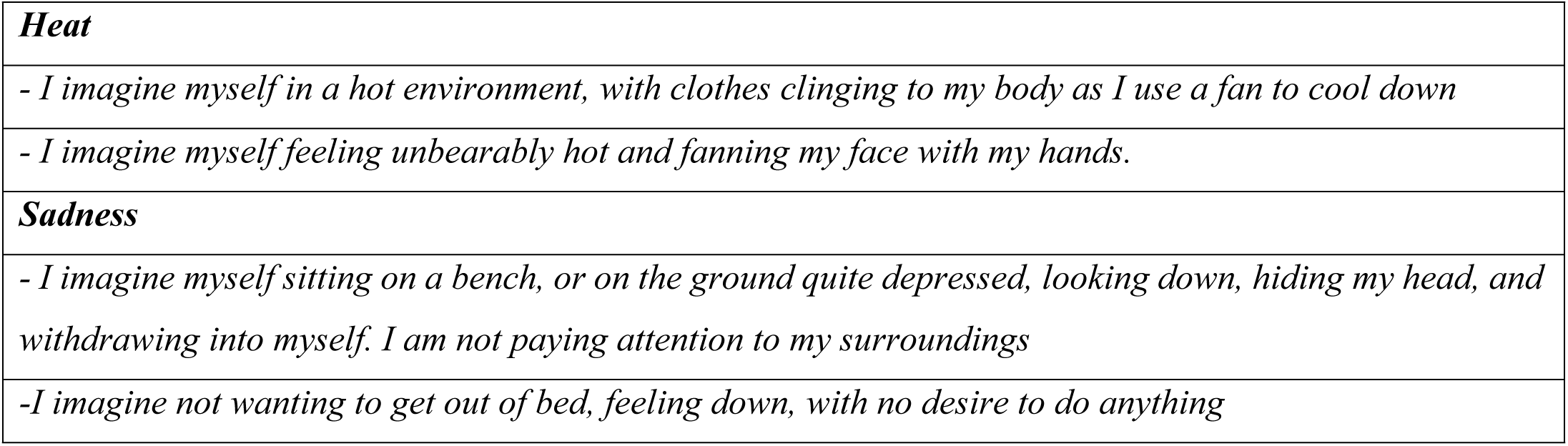
Examples of suggested exercises for imagery training.

|  |
| --- |
| ***Movement*** |
| - *I imagine myself running freely on the soccer field with my teammates* |
| - *I imagine being in a tight crowd and trying to move through it* |
| ***Thirst*** |
| - *I imagine myself dehydrated with a dry mouth, yearning for an icy bottle of water on a sweltering day* |
| - *I go to the vending machine, grab a bottle of refreshing sparkling water, and finish it immediately* |
| **Heat** |
| - I imagine myself in a hot environment, with clothes clinging to my body as I use a fan to cool down |
| - I imagine myself feeling unbearably hot and fanning my face with my hands. |
| **Sadness** |
| - I imagine myself sitting on a bench, or on the ground quite depressed, looking down, hiding my head, and withdrawing into myself. I am not paying attention to my surroundings |
| - I imagine not wanting to get out of bed, feeling down, with no desire to do anything |

The experiment took place inside an electromagnetically and acoustically shielded booth (Faraday cage). Participants were seated in a comfortable chair positioned 114 cm away from a screen located outside the booth and wore *Trust*-brand headphones to listen to the audio stimuli. During the recording session, participants were instructed to fixate on a central point—a glowing cross positioned at the center of the screen—while refraining from moving their eyes, blinking, swallowing, or making any other movements, all while maintaining a relaxed posture.

The experimental task involved listening to the auditory cues and recalling the corresponding motivational state, as vividly and realistically as possible, as if experienced in the first person. During the imagery task, a yellow frame appeared on the screen to which ERP signals were synchronized. At the end of each sequence, the participant was presented with a question related to the stimulation content (e.g., Has it been mentioned (in this experimental sequence) to desire a nice plate of pasta with meat sauce?) in the center of the screen and instructed to answer “yes” or “no” verbally a few seconds later. The question served to verify the participant’s actual execution of the task and attentional engagement.

Each stimulus sequence began with three warning signals (“ATTENTION,” “READY,” and “GO”), displayed for 1 second in capital letters, and ended with “THANK YOU!” The words appeared in white on a gray background, in Times font, approximately 3 cm in size. Sequence presentation was randomized across participants, with consistent screen brightness and audio volume. Luminance conditions were scotopic. Each audio stimulus lasted 2500 ms, followed by a brief 100 ± 20 ms interval (ISI), after which a yellow frame appeared for 2500 ms. A 60 ± 20 ms interval (ITI) preceded the next audio stimulus (see Fig. 1). Each item, consisting of audio and imagery task, lasted 5160 ± 40 ms. Overall, each sequence lasted about 3 minutes, with brief pauses, and the entire experiment took approximately 1.5 hours. A total of 18 sequences were presented, preceded by a shorter practice sequence. After the EEG recording, participants completed a Likert-scale questionnaire (1 = very difficult, 5 = very easy) assessing the ease of recall of motivational states related to primary needs, physical sensations, secondary needs, and emotions. The stimuli were presented using *Eevoke v2.2* (ANT Neuro).

**Fig. 1.**
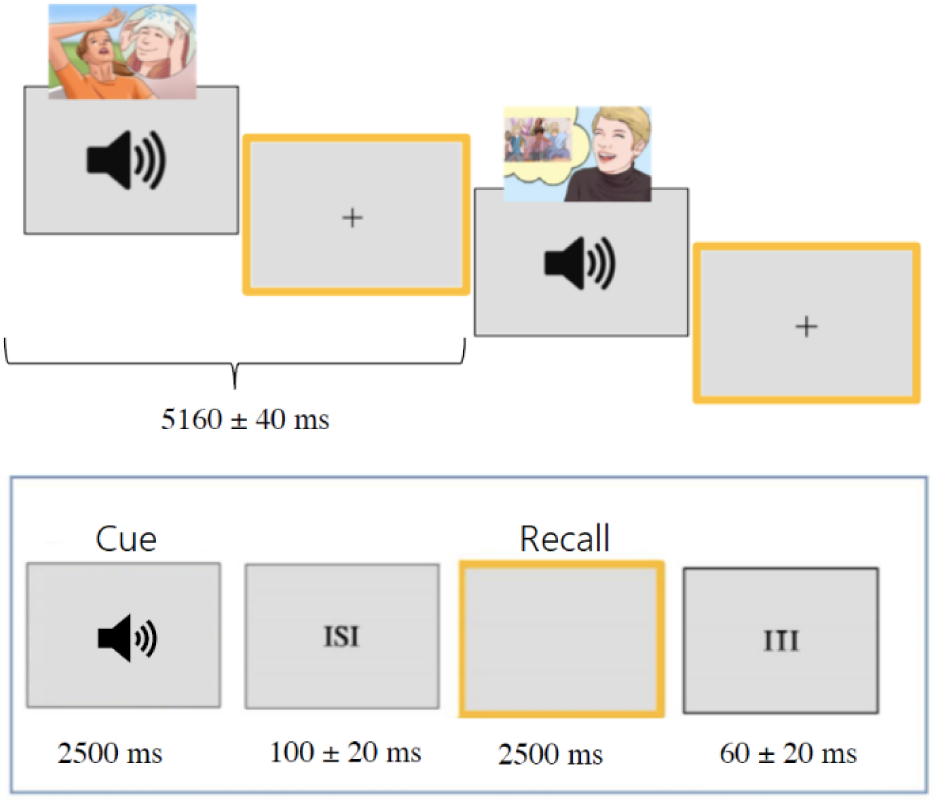
Timeline of the experimental procedure.

### EEG recording and analysis

Brain activity was monitored using 128 electrodes placed according to the 10-5 International system, with horizontal and vertical electro-oculograms also being recorded. The averaged mastoids were used as the reference. Electrode impedance was kept below 5 kΩ, and the sampling rate was 512 Hz. EEG and EOG signals were captured using the *Cognitrace* system (ANT Software) and amplified with a bandpass filter (0.16–70 Hz). Any EEG artifacts exceeding ±50 μV were automatically rejected before averaging. EEG epochs synchronized with stimulus presentation were processed through the *EEProbe* system (ANT Software). ERPs were averaged offline from 100 ms before to 1200 ms after stimulus onset (of prompt for imagery).

The mean area amplitude of the anterior N400 component, measured between 400–600 ms at electrodes AF3, AF4, F7, and F8, and of the centroparietal P400, measured between 400–600 ms at electrodes CP1, CP2, and CPZ, was calculated. The choice of the electrode and the time window was based on where and when the components reached their maximum amplitude on the scalp, and on the previous literature. Measurements were conducted on individual ERP signals averaged across the 12 micro-categories and 4 macro-categories during the imagery task. The mean area amplitudes of the various components were subjected to repeated-measures ANOVA, with the following factors: motivational state (4 levels in the macro-category analysis and 12 levels in the micro-category analysis), electrode (depending on the component), and hemisphere (left, right). Using ASA software (ANT Software, Enschede, The Netherlands), characteristic topographical maps for the 4 macro-categories were generated, enabling visualization of the spatial distribution of the primary ERP components identified in the study: N400 and P400. These maps were created by plotting isopotential lines onto a color scale, obtained by interpolating voltage values between surface electrode sites. Overall. N400 component exhibited larger amplitudes at anterior regions, while P400 component was more pronounced at centroparietal sites.

### Source reconstruction

To identify the cortical sources of surface electrical activity in the P/N400 time window four swLORETA models were conducted on mean ERP averages corresponding to each macro-category of motivational state. *Low-Resolution Electromagnetic Tomography* (LORETA) is a powerful source reconstruction technique able to localize neural activity with high spatial resolution by estimating the sources of electrical activity within the brain (Pascual-Marqui, 2011). In this study we used and advanced algorithm proposed by Palmero-Soler et al. (2007) called SwLORETA, which incorporates a *Singular Value Decomposition* (SVD) based lead field weighting. Additionally, synchronization tomography and coherence tomography based on SwLORETA were introduced to analyze phase synchronization and standard linear coherence, applied to current source density.

## Results

### Self-reported data

#### Ease of recall

Data were collected from the whole sample of 30 students (13 male, 17 female), with an average age of 22.4 years (SE = 1.98). Mean scores were subjected to a one-way ANOVA featuring factor motivational state (4 levels of variability). Results indicated significant differences (F (3,87) = 8.25; p < 0.05); post-hoc comparisons revealed that “visceral needs” (i.e., hunger, thirst and sleep) were significantly more easily imagined (M = 4.40; SE = 0.10) than other motivational states: “somatosensory” (M = 3.73; SE = 0.18) with p < .05; “secondary” (M = 3.47; SE = 0.16) with p < .001, and “affective” (M = 3.33; SE = 0.24) with p < .001.

#### Stimulation content

The percentage of correct responses to the final test (i.e. the final question about the run content) was 91.90% (SE = 7.04), corresponding to approximately 1.52 errors out of 18 questions (SE = 1.24). This indicates that participants paid close attention to the auditory stimulus semantic content as required for the imaginative task.

### Electrophysiological results

#### Anterior N400 component (400-600 ms)

##### Macro-categories

The ANOVA performed on N400 mean area amplitude values recorded at AF3, AF4, F7, F8 sites in the 400-600 ms time window showed the significance of “state” factor (F (3, 66) = 3.10; p <0.05), with larger N400 amplitudes during “affective” (M = −0.40; SE = 0.67) than “visceral” (M = 0.23; SE = 0.51) (p<0.05) and “secondary” states (M = 0.21; SE = 0.58) (p <0.01) as can be appreciated from ERP waveforms of Fig. 2 and topographical maps displayed in Fig. 3. The significant interaction of “state” x “electrode” (F (3,66) = 2.69; <0.05) and relative post-hoc comparisons showed that N400 was larger (p <0.01) to “somatosensory” (M = −0.48; SE = 0.51) than “visceral” (M = 0.22; SE = 0.40) and “secondary” states (M = 0.26; SE = 0.44) at anterior frontal sites. The largest N400 response (p <0.01) was recorded during recall of “affective” states at inferior frontal sites (M= −0.432, SE= 0.54),

**Fig. 2.**
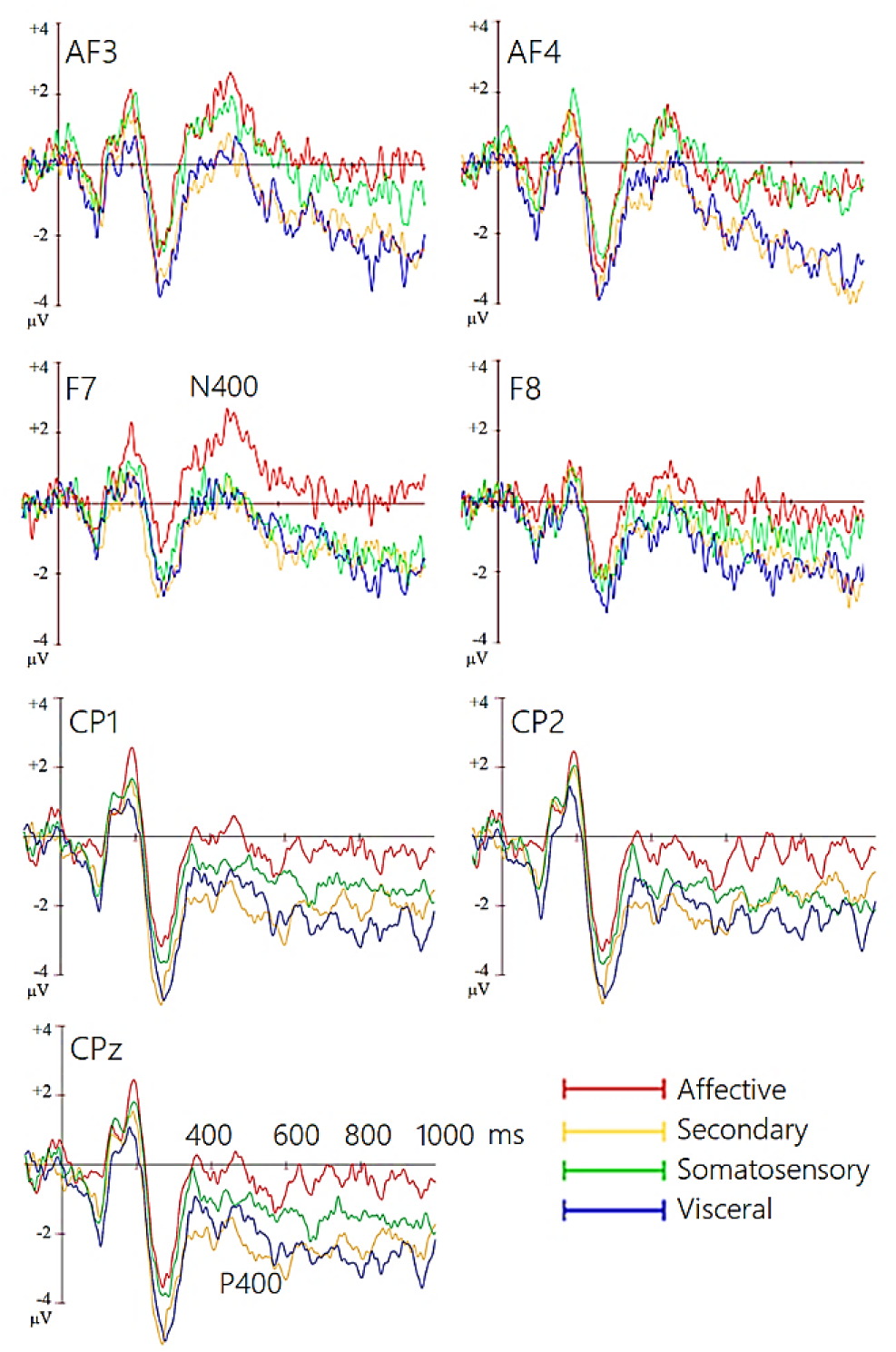
Grand-average ERP waveforms recorded from left and right anterior and inferior frontal (top) and left, right and midline centroparietal sites (bottom) during recall of the 4 macro-categories of motivational states.

**Fig. 3.**
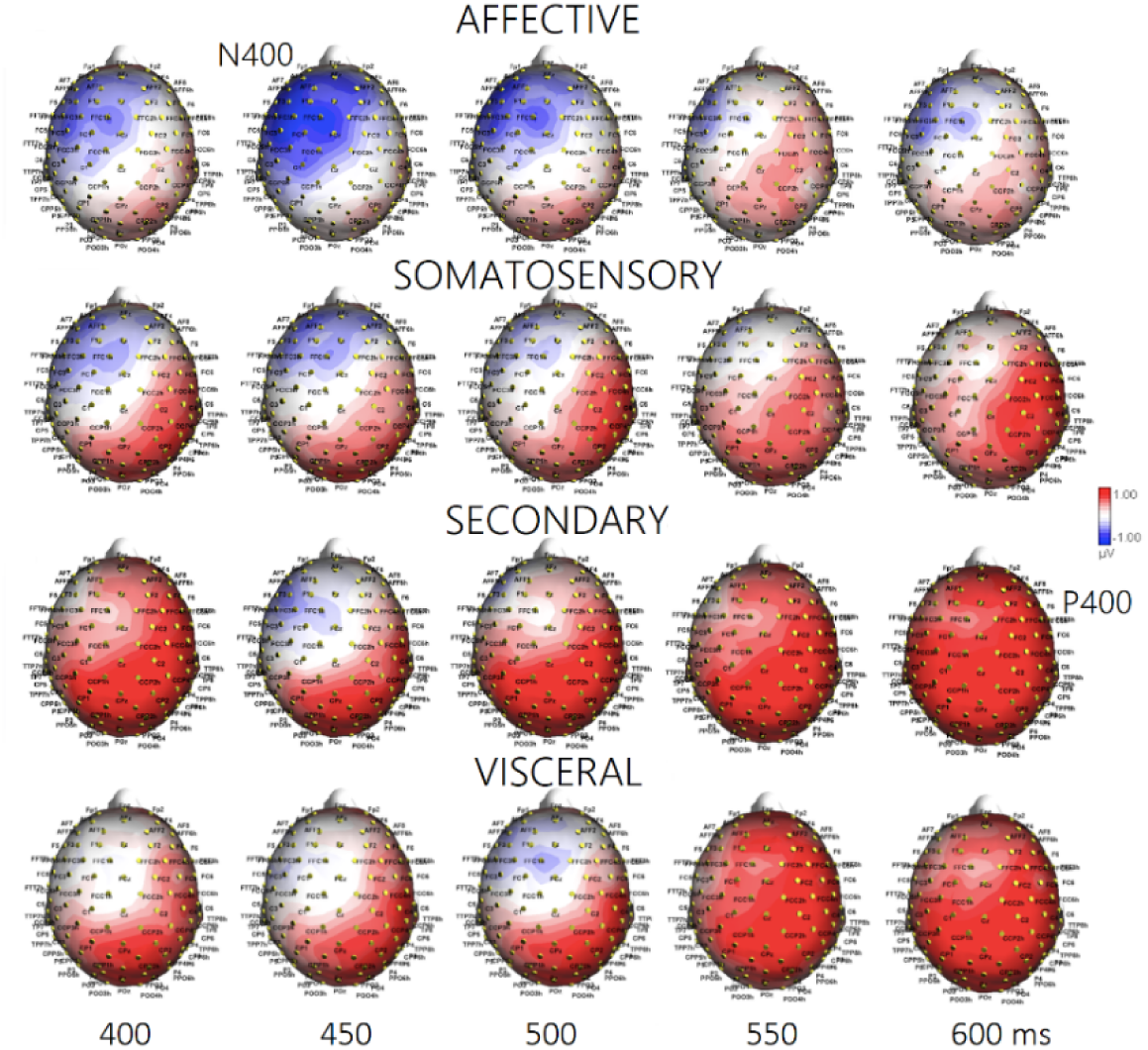
Isocolour topographical maps of voltage potentials recorded during recall of the 4 macro-categories of motivational states in the 400-600 ms time window (50 ms step).

##### Micro-categories

The ANOVA performed on N400 mean area amplitude values recorded in association with microstates yielded the significance of “state” factor (*F* (11, 242) =2.25; *p<.05*). Significances at post-hoc comparisons, along with N400 mean amplitude values for each of the microstates, and standard deviations, are reported in Fig 4.

**Fig. 4.**
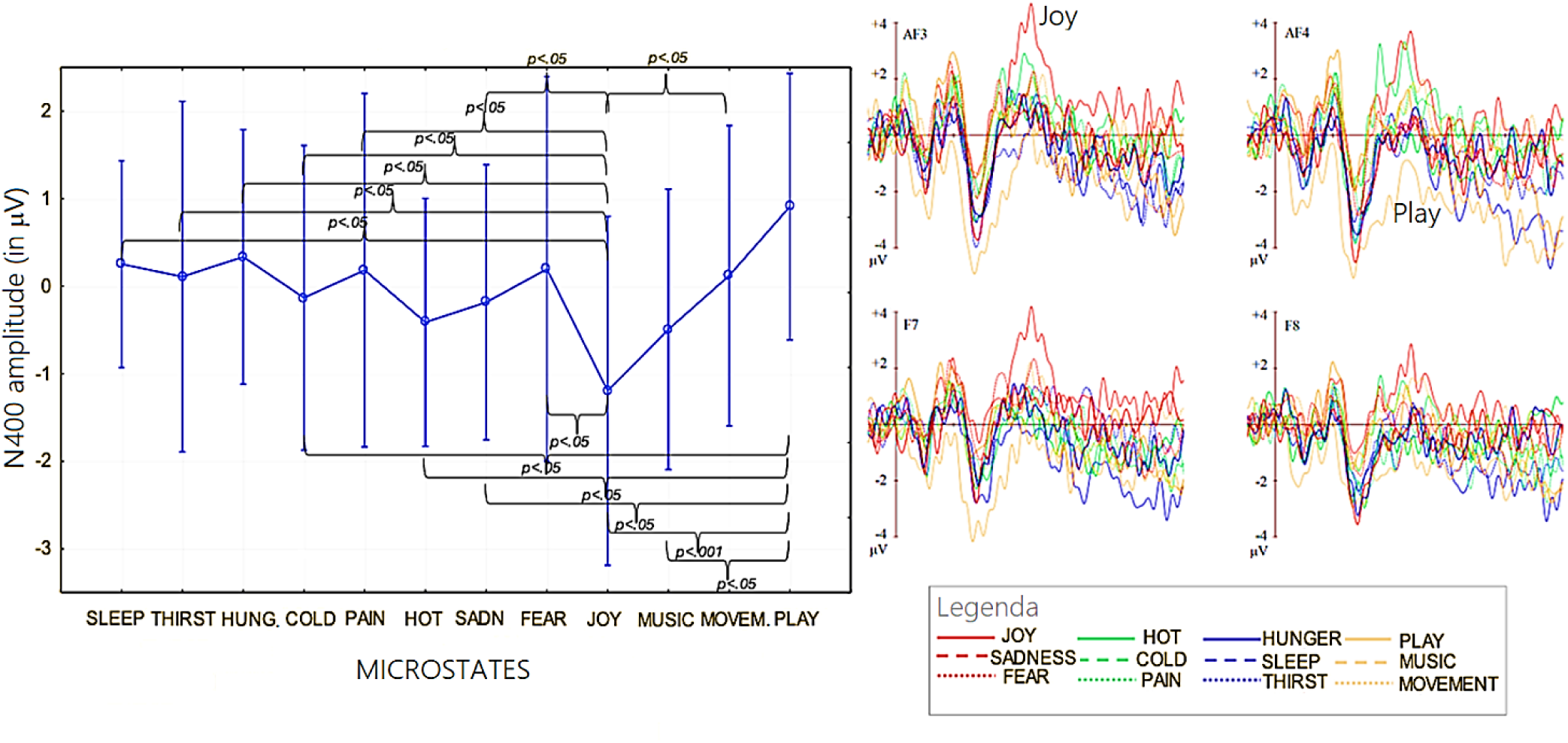
(Left) Mean area amplitude of N400 response recorded as a function of the 12 micro-categories of motivational states. Standard deviations and statistical significances at post-hoc comparisons are also shown for the various states. (Right) Grand-average ERP waveforms recorded from left and right anterior and inferior frontal sites during recall of the 12 microstates.

Post hoc tests indicated that “joy” was the motivational state marked by the highest N400, in contrast to “social play,” which was characterized by the lowest N400 amplitude. Figure 5 (left) shows the grand-average ERPs recorded in association with the 12 micro-categories at anterior and inferior left and right frontal electrodes, and at lateral and midline centroparietal electrodes (right), separately for each reference microstate.

**Fig. 5.**
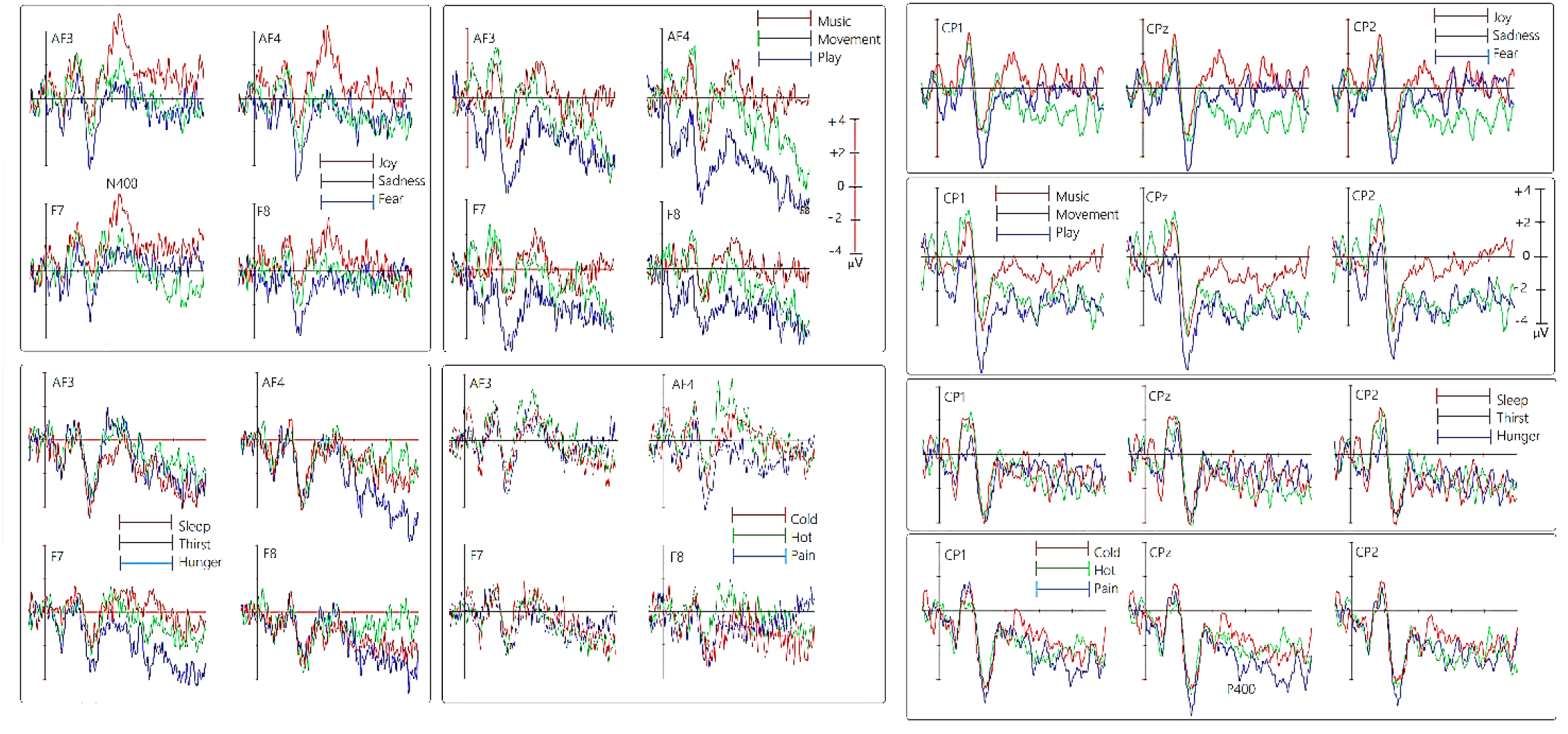
Grand-average ERP waveforms recorded from left and right anterior and inferior frontal (left) and left, right and midline centroparietal sites (right) during recall of the 12 micro-categories of motivational states.

The study also examined whether a correlation existed between the ease of recalling motivational states, expressed in Likert scores ranging from 1 to 5, and the mean area amplitude of N400 responses recorded during each microstate.

A Spearman’s Rho correlation test was conducted on the two data sets, showing a linear correlation: the greater the difficulty in recalling a state, the larger the N400 amplitude (r = 0.50612; p < 0.05), as can be appreciated in Fig. 6. This finding likely suggests that the N400 component reflected the cognitive effort required to evoke or recall motivational states.

**Fig. 6.**
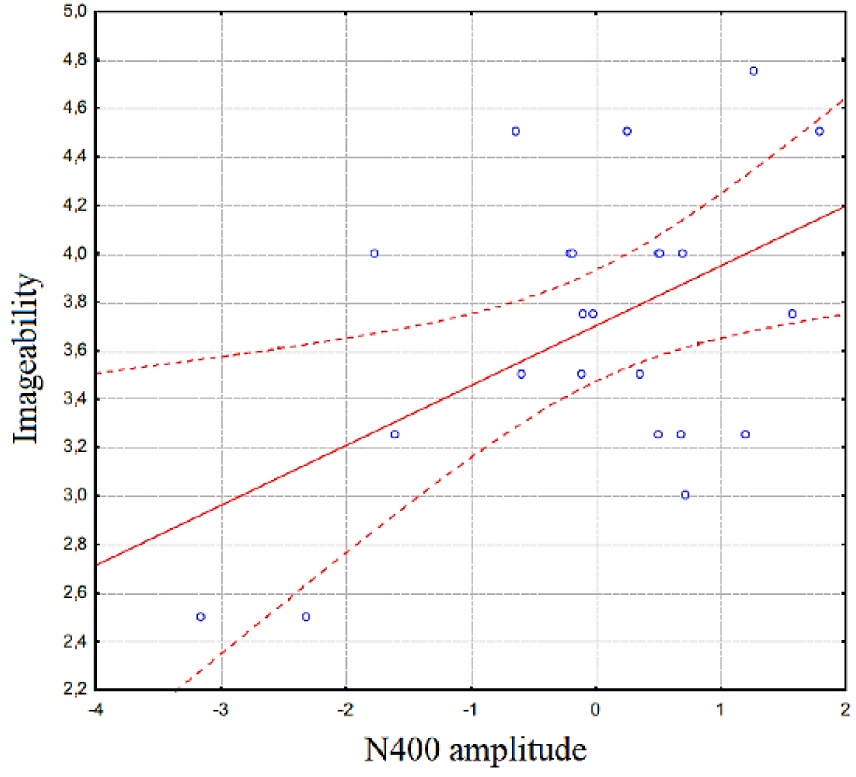
Spearman Rho correlation between subjective rates of complexity of recalling the motivational states and individual N400 mean amplitude values.

#### Centroparietal P400 response

##### Macrocategories

The ANOVA performed on P400 mean area amplitude values recorded at CP1, CP2 e CPZ sites in the 400-600 ms time window showed the significance of “state” factor (F (3,66) = 2.84; p<0.05): post-hoc comparisons showed larger P400 responses during recall of “secondary” (M = 1.13; SE = 0.61) (p <0.008) and “visceral” (M = 0.92; SE = 0.75) (p <0.03) than “affective” states (M = 0.23; SE = 0.59), with intermediate amplitude during recall of “somatosensory” states (M=.60; SE=.74), see Fig. 2 (Lower) and 3 for ERP waveforms and topographical maps.

##### Microcategories

The ANOVA performed on P400 amplitude values recorded in association with microstates yielded the significance of “state” factor (*F* (11, 242) =2.05; p<0.025). Significances at post-hoc comparisons, along with P400 mean amplitude values for each of the microstates, and standard deviations, are reported in Fig 7.

**Fig. 7.**
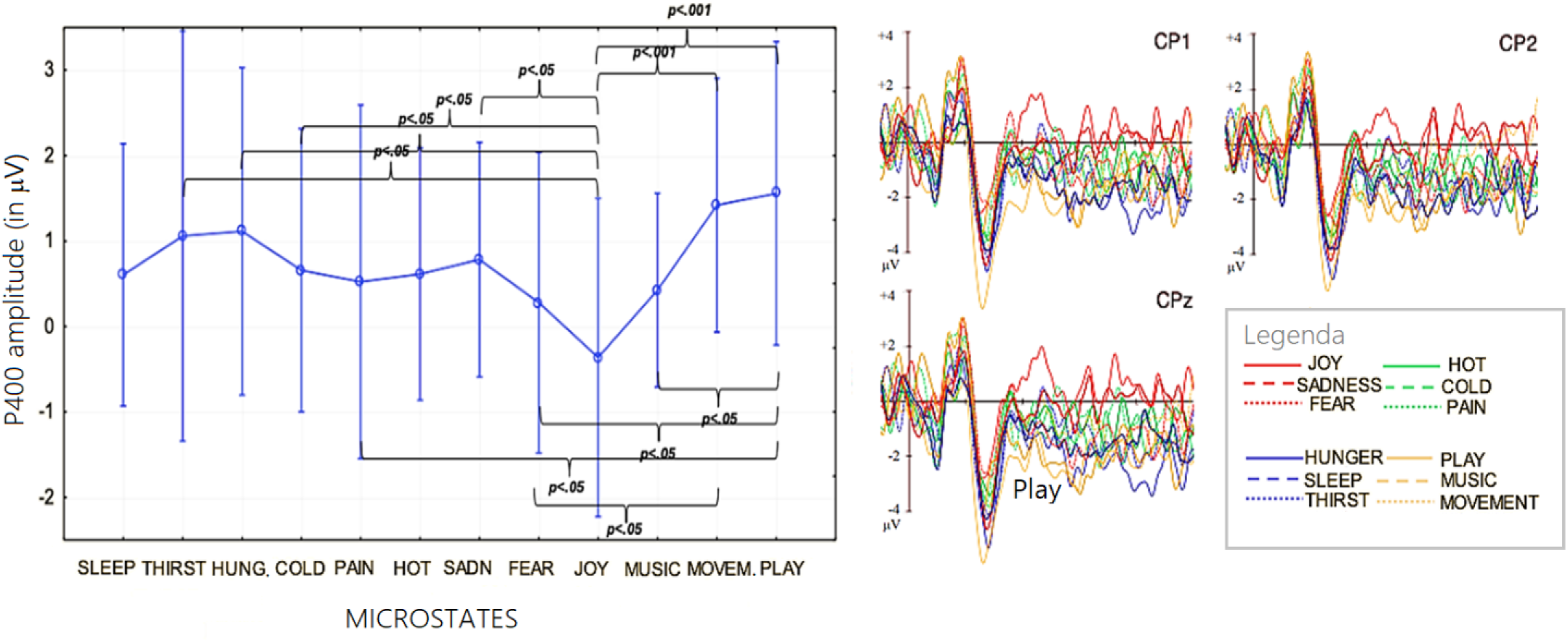
(Left) Mean area amplitude of P400 response recorded as a function of the 12 micro-categories of motivational states. Standard deviations and statistical significances at post-hoc comparisons are also shown for the various states. (Right) Grand-average ERP waveforms recorded from left, right and midline centroparietal sites during recall of the 12 microstates.

Post hoc tests indicated that “play” (p<0.0002) and “movement” (p<0.0006) were the motivational state marked by the highest P400, in contrast to “joy” and “fear” which were characterized by the lowest P400 amplitude. Figure 5 (right) shows the grand-average ERPs recorded in association with the 12 micro-categories at anterior and inferior left and right frontal electrodes, and at lateral and midline centroparietal electrodes, separately for each reference microstate. It can be concluded, therefore, that the amplitude of the ERP component P400 recorded at centro-parietal sites changed depending on the mentally represented micro-category.

The correlation between P400 mean amplitude values and the ease in recalling motivational states was not significant. This suggests that imaginative effort or efficacy was expressed more by N400 responses.

#### swLORETA source reconstruction

In Table 2 are listed the active electromagnetic dipoles significantly explaining surface potentials within the 400-600 ms time window during the recall of the four motivational states according to swLORETA (Palmero-Soler et al., 2007). Almost all macrostates featured the activation of the left superior frontal gyrus (BA10) involved in Self-representation and working memory (inner voice), as well as posterior brain regions involved in Imagery.

**Table 2.**
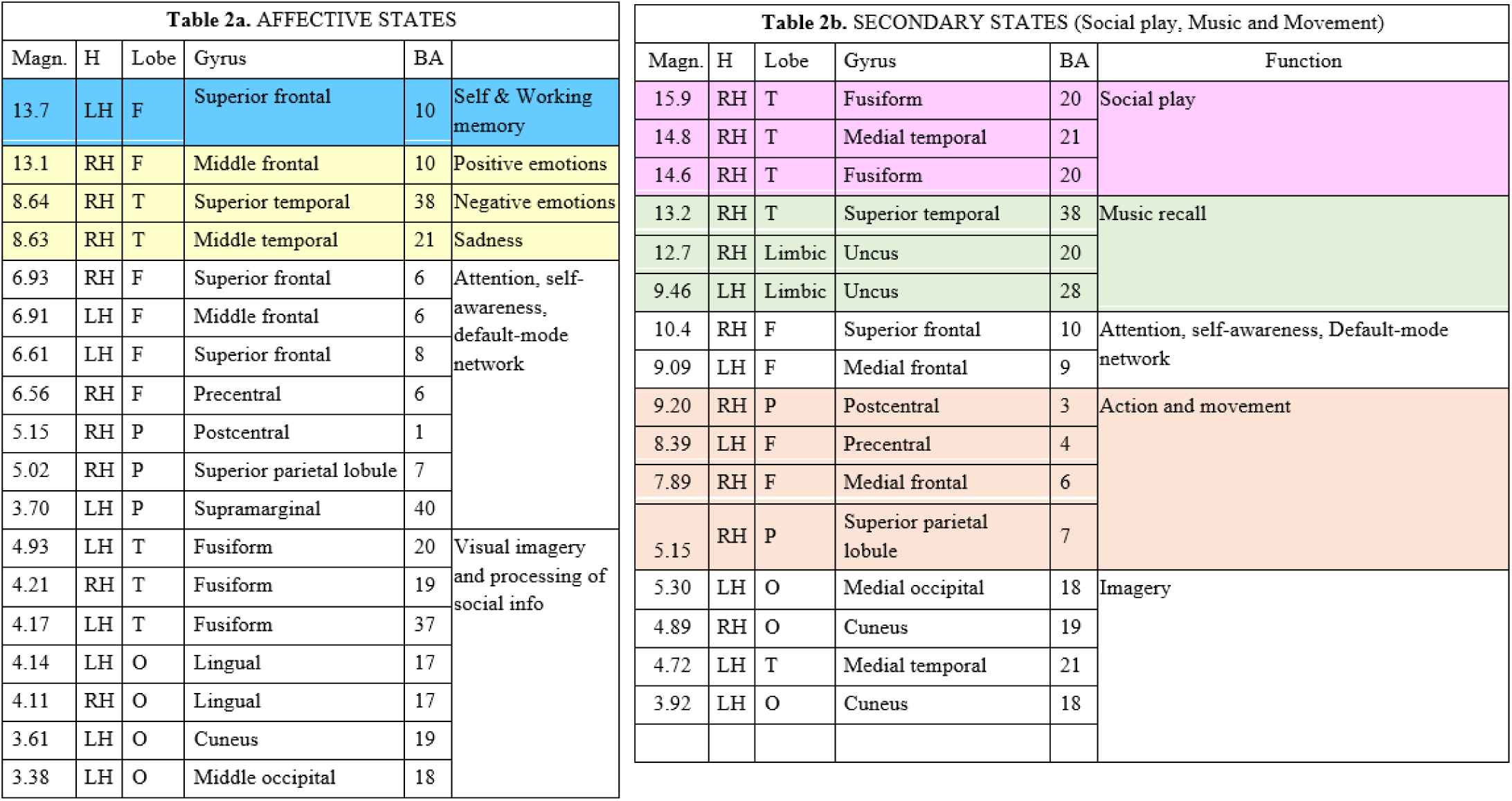
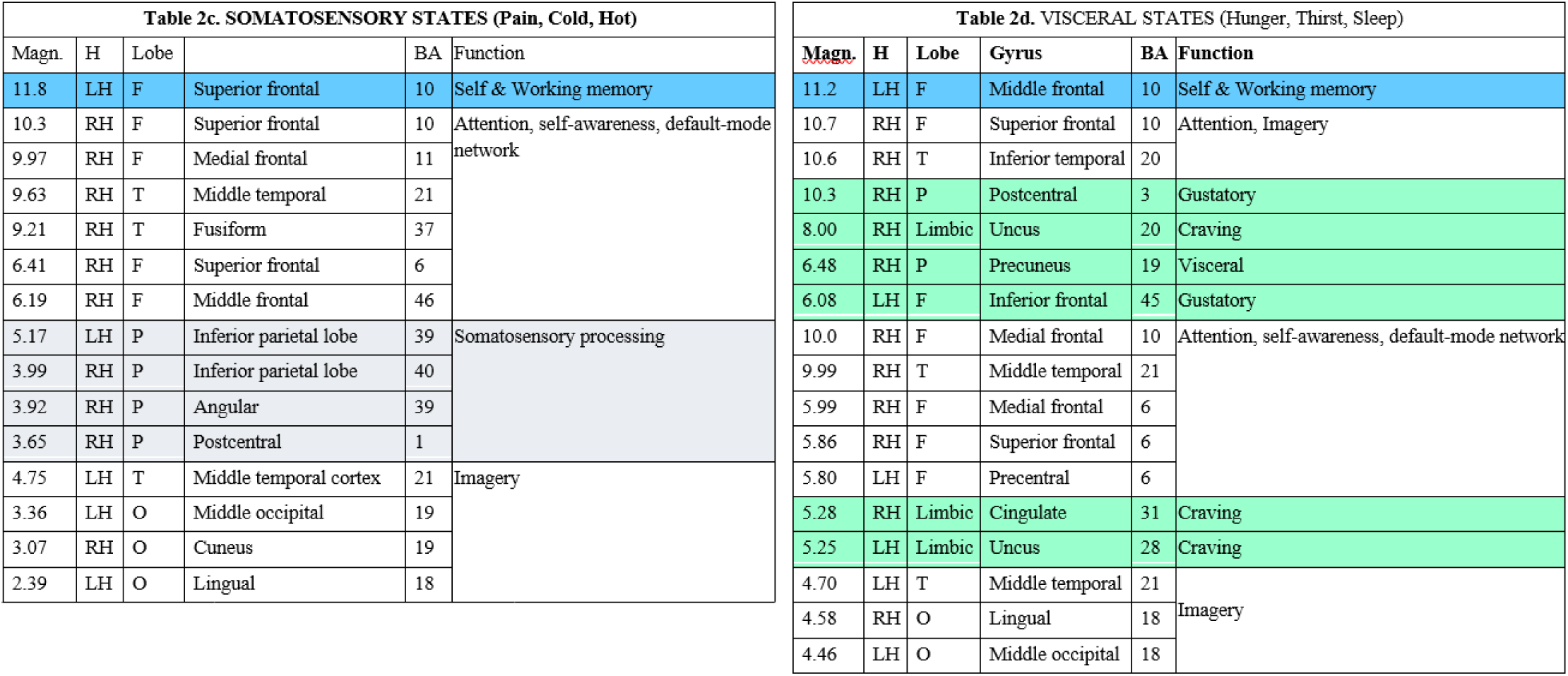
List of active electromagnetic dipoles (along with their Talairach coordinates) explaining brain voltage during the four recalled macromotivational and physiological states (group analysis). Magn. = magnitude in nA; H = Hemisphere, BA= Brodmann areas, function= presumed functional properties.

Affective motivational states (Table 2a) specifically involved right fronto-temporal areas, namely the right medial frontal (MFG), superior temporal, and middle temporal gyri (MTG). Additionally, several other neural areas traditionally associated with visual imagery were activated, including the fusiform, lingual, and supramarginal gyri. Fig. 8 displays the localization of more active areas as projected in sagittal, coronal and axial MRI sections.

**Fig. 8.**
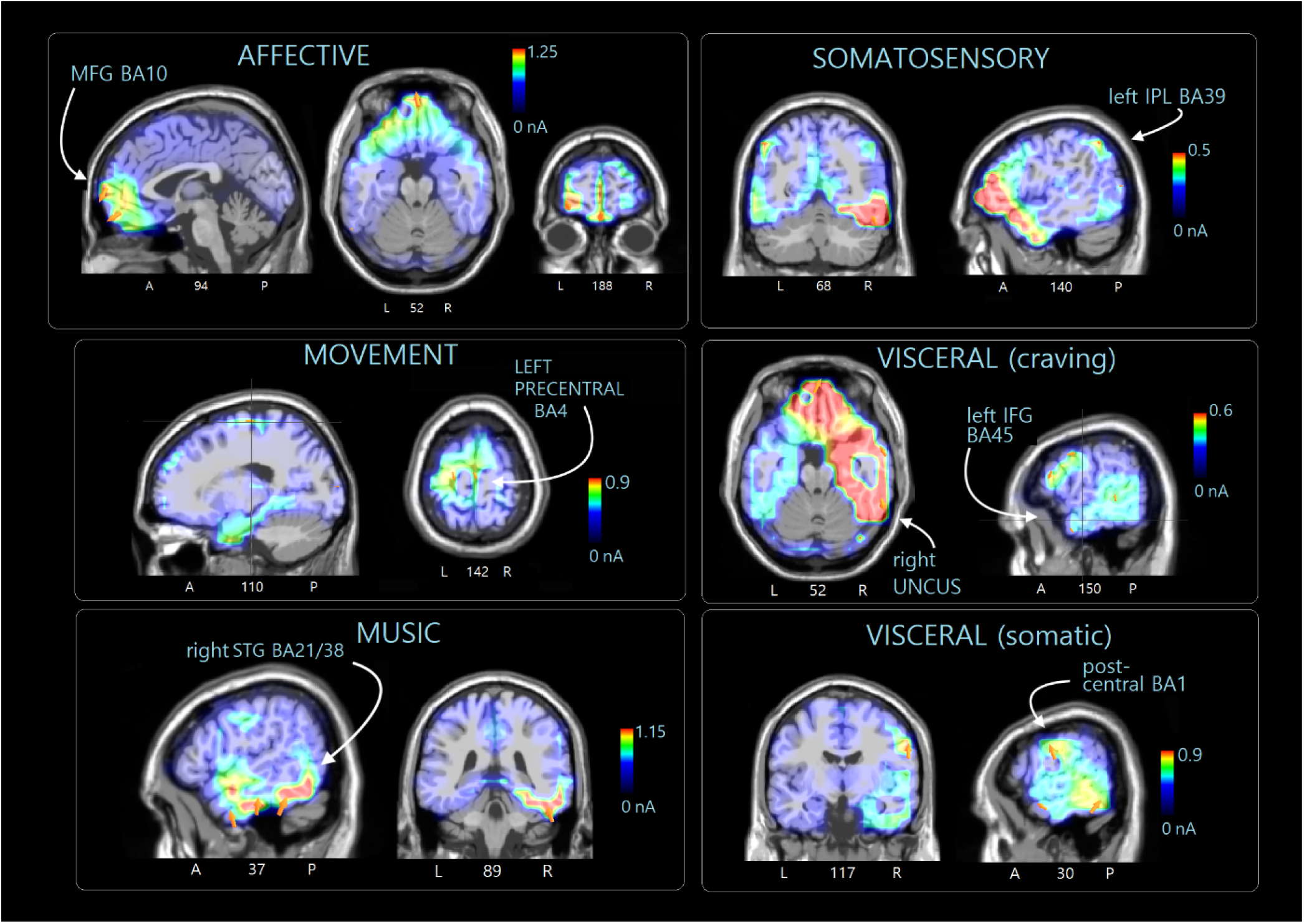
Sagittal, axial and coronal views of swLORETA source reconstructions of N400 surface potentials recorded in the 400-600 ms time window during the various macro-categories of motivational states. The various colors represent differences in the magnitude of the electromagnetic signal (nA). The electromagnetic dipoles appear as arrows and indicate the position, orientation and magnitude of the dipole modelling solution applied to the ERP waveform in the specific time window. A, anterior; P, posterior; L, left; R, right; numbers refer to the displayed brain slice in the MRI imaging plane.

Table 2b lists the more active dipoles during the recollection of “secondary” desires, which included the right MTG, STG (BA38) and the bilateral limbic areas (BA20 and Ba28) possibly linked to musical imagery, and left precentral gyrus (BA4) linked to motor imagery (see neuroimages in Figure 8).

In Table 2c, the areas involved during the recall of “somatosensory” sensations are listed. The most distinctive areas appeared to be the angular (BA39) and postcentral gyri on the right (BA1), as well as the left and right inferior parietal lobes (BA40) (see Figure 8). Table 2d lists the areas involved in the subjective recall of visceral needs. The most active areas included the right MFG and IFG, right precuneus, left and right uncus involved in craving and hunger, besides imagery related areas (e.g., right lingual gyrus and left middle occipital gyrus), as shown in Fig. 8.

#### ROI analysis

In order to have a comprehensive view of the specific sources of electro-magnetic activity across the 4 motivational states, 8 regions of interest (ROIs) per hemisphere were identified following the ROI clustering procedure used to perform statistical analyses on individual LORETA solutions (Babiloni et al., 2004, 2006; Cannon et al., 2008, 2009). The selected ROIs are listed in Table 3.

**Table 3.** List of Regions of Interest identified and referenced to the Gyri and Brodmann areas (BAs) included in each cluster. A similar clustering was used in Della Vedova and Proverbio (2024) and in Proverbio and Cesati (2024).

| ROIs | BA | GYRUS |
| --- | --- | --- |
| OCC | 18, 19 | OCCIPITAL CORTEX: Inferior Occipital, Middle Occipital, Superior Occipital, Lingual (also BA 17) gyri, Cuneus (also BA 17) |
| FUSIF | 19, 20, 37 | FUSIFORM AREA: Fusiform Gyrus, Cerebellum (Anterior Lobe), Posterior lobe (Declive), Middle occipital (only BA 37) and Inferior temporal gyri (only BA 37) |
| TEMP | 19, 20, 21, 22, 38, 39, 41, 42 | SUPERIOR, MIDDLE AND INFERIOR TEMPORAL CORTEX: Superior Temporal, Middle Temporal and Inferior Temporal Gyri |
| PREM | 4, 6 | PREMOTOR CORTEX: Precentral (also BA 43), Middle Frontal, Medial Frontal, Superior Frontal, Precentral gyri, Paracentral Lobule |
| FRONTAL | 8, 9, 46 | MEDIAL, SUPERIOR FRONTAL AND DORSOLATERAL PREFRONTAL CORTEX: Middle Frontal, Medial Frontal and Superior Frontal Gyri |
| ORBIF | 10, 11, 44, 45, 47 | ORBITOFRONTAL AND INFERIOR FRONTAL CORTEX: Superior Frontal, Medial Frontal, Middle Frontal, Inferior Frontal and Rectal Gyri |
| PARIETAL | 1, 2, 3, 7, 19, 39, 40 | PARIETAL CORTEX: Superior and Inferior Parietal Lobule, Precuneus, Postcentral, Supramarginal and Angular Gyri |
| LIMBIC | 20, 23, 24, 28, 31, 34, 35, 36, 38 | LIMBIC AREA: Uncus, Cingulate Gyrus, Anterior Cingulate, PPA |

The source reconstruction data enabled the identification of the most active Regions of Interest (ROIs, in nA) during the recall of various motivational macrostates. Figure 9 visually depicts the most engaged ROIs in the right and left hemispheres (in terms of total strength of active ROIs), highlighting a distinct pattern that lends itself well to potential classification using deep learning techniques. A 2 way ANOVA applied to average magnitudes computed as a function of hemisphere and ROI (as within factors) and macro-category (as between factor) yielded the significance of Hemisphere factor (F1, 28= 8.97, p <0.006), with larger signals over the right (RH = 11.01; SE= 1.83) than left (LH = 5.03, SE= 0.87 μV) hemisphere. The further interaction of Hemisphere x ROI (F7,24 = 2.74; p <0.03) and relative post-hoc comparisons showed stronger right-sided signals especially over temporal, fusiform and orbitofrontal ROIs.

**Fig. 9.**
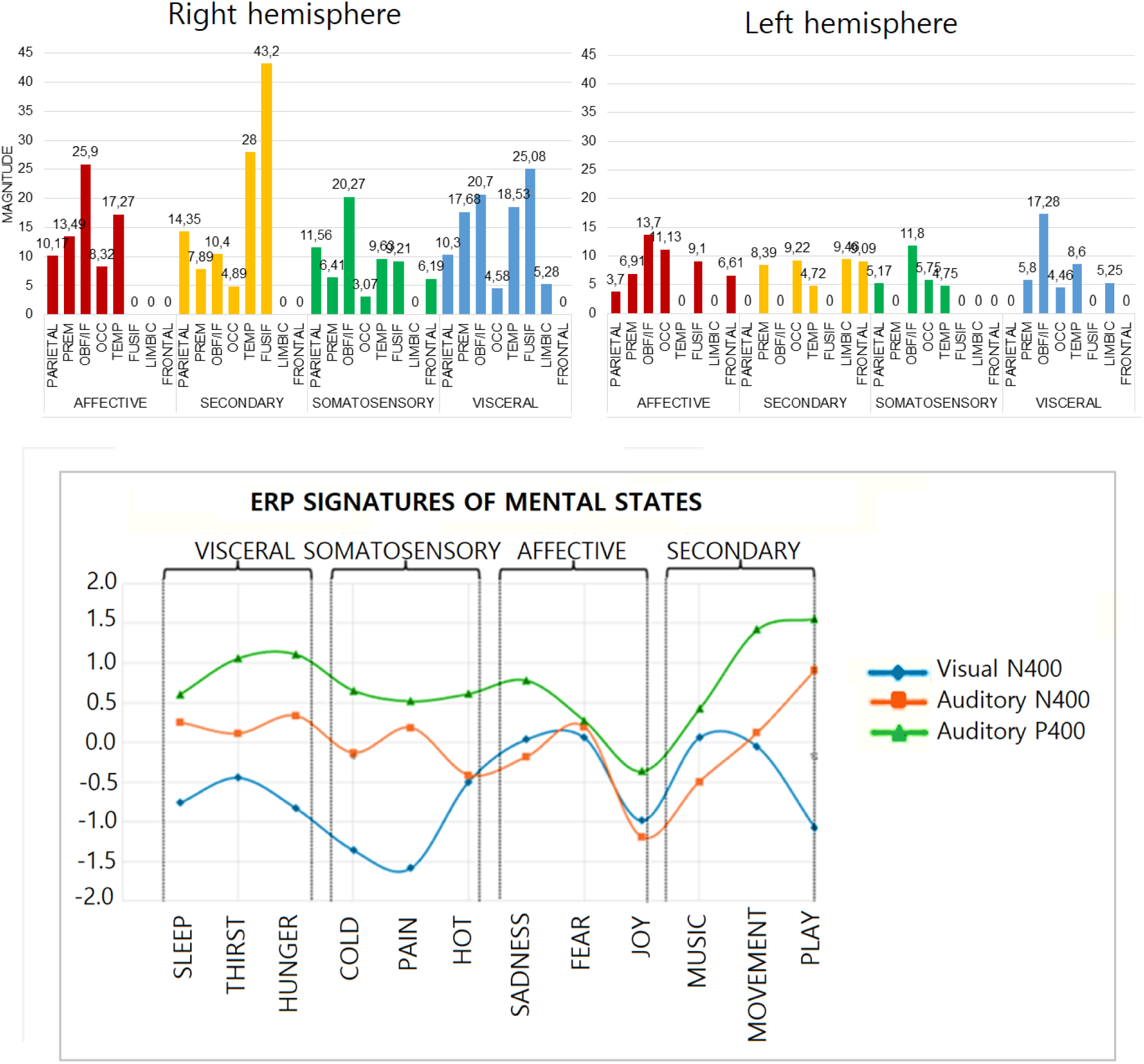
(Top) Cumulative values of average magnitude (dipole strength in nA) of electromagnetic dipoles found active in each of the selected ROIs of the left and right hemisphere, during the recall of states belonging to the 4 macro-motivational states. (Bottom) The amplitude trend profile of the P/N400 components recorded in the present, and in Proverbio & Pischedda’s (2024) studies, based on visual pictograms, shows a clear similarity in the behavior of these neural signatures, irrespective of the experimental condition.

## Discussion

This study investigated ERP correlates associated with 12 motivational states elicited through mental representations, organized into four macro-categories. Significant differences emerged across both micro-and macro-categories, affecting late latency components (N400 and P400). A for previous studies with a similar paradigms (e.g. Proverbio et al., 2022, 2023), no category-specific effects on sensory or perceptual components (i.e., N170, N2, P2, or P300) was observed during recall of imagined states. Indeed, early ERP overlaps reflect processing of the identical bright frame (the visual cue with which ERP epochs were synchronized), with no differences attributed to category-related imaginative processes earlier than 300 ms (Dijkstra et al., 2018; 2020). Similar to the ERP study in which motivational states were prompted using pictograms (Proverbio and Pischedda, 2023), here an anteriorly negative N400 component with a 400-600 ms latency, as well as a centro-parietal positive P400 within the same time window, were identified. However, unlike the current study’s auditory cues, the pictogram-based recall elicited a later positive component (LPP with an 800-1000 ms latency), likely due to the additional processing time required to interpret visual pictograms compared to auditory cues (Shelton and Kumar, 2010). In both studies, the N400 component localized primarily to anterior frontal and inferior frontal regions, suggesting a common neural mechanism for mentally activating motivational states. Frontal and fronto-temporal areas are known to play a pivotal role in imaginative processes, specifically in initiating, controlling, and sustaining mental representations (Spagna et al., 2021; Dijkstra et al., 2017). The N400 amplitude was notably greater when recalling “affective” states, particularly joy, as well as “somatosensory” states, such as pain and cold, paralleling findings from the previous pictogram-based study. Overall, brain activity in the P/N400 latency stage was larger over the right hemisphere, particularly for over temporal, fusiform and orbitofrontal ROIs: this finding fits with previous data by Proverbio et al. (2023) that similarly demonstrated increased right hemisphere involvement during imagery tasks, particularly for the LPP component, and other literature on mental imagery (Ehrlichman and Barrett, 1983; Gasparini et al., 2008; Liu et al., 2022).

The N400 component appears to be highly sensitive to cognitive effort related to motivational states. Conceptually, larger N400 amplitudes, particularly in anterior regions, reflect greater cognitive processing when recalling or distinguishing states with high emotional salience, such as “affective” states, indicating an increased mental load associated with emotionally charged memories. For micro-categories, N400 variations suggest that distinct motivational nuances (e.g., “joy” versus “social play”) require different degrees of cognitive engagement, with heightened N400 amplitudes linked to more challenging or intense states. The correlation between N400 amplitude and the subjective difficulty of recall further supports the role of N400 as an index of cognitive effort in retrieving complex or personally significant motivational experiences. The P400 component, more prominent at centroparietal sites, reflects categorical distinctions in motivational representation, particularly among different types of internal states (e.g., “secondary” or “visceral” versus “affective” states). While less sensitive to recall difficulty, P400 amplitudes vary with specific micro-categories, indicating that this component may capture differences in the sensory or embodied aspects of motivational states, particularly for states involving movement and play, without directly indexing cognitive effort.

In sum, the N400 aligns with cognitive effort linked to emotional recall, while the P400 provides insight into the differentiation of motivational states based on their embodied characteristics. Overall the amplitude of the ERP component anterior N400 (see also Dijkstra et al., 2020) and centro-parietal P400 significantly changed depending on the mentally represented micro-category. In this study source reconstruction of scalp-recorded potential was also carried out for “mind reading” purposes.

The swLORETA reconstruction of intracortical generators has identified consistent brain activations across several macrocategories, uniquely associated with the imagery task. Specifically, the left superior frontal gyrus demonstrated significant activation in “affective” and “visceral” motivational states, while the left middle frontal gyrus was more prominently activated during “somatosensory” states. These regions appear pivotal for processes of imagination and working memory, as suggested by Spagna et al. (2024), where the left superior frontal gyrus showed heightened activation during visual imagination tasks in comparison to control conditions. Similarly, Lima et al. (2015) underscores the left middle frontal gyrus’s role, noting its strong correlation with the vividness of auditory imagery. Furthermore, scientific evidence indicates that electrical stimulation of these regions can induce complex visual hallucinations (Blank et al., 2000). The left superior frontal gyrus also acts as a key area for self-representation (the “inner voice,” as discussed in González et al., 2018) and self-referential processing (Depalma and Proverbio, 2024; D’Argembeau et al., 2007), which are essential for the internal representation of personal needs. According to Skottnik and Linden, 2019), the frontoparietal network additionally supports episodic memory, while Barry et al. (2019) highlight the role of the ventromedial prefrontal cortex (vmPFC) in mental representation. Patients with vmPFC damage often experience difficulties in imagining scenes unless prompted by highly specific cues, suggesting that the vmPFC plays a crucial role in selecting the elements appropriate to a specific scene. The occipitotemporal cortex (BA18, 19, 21, 37) remained consistently active across all imagery conditions, aligning with findings from visuo-spatial imagery studies (e.g., Bartolomeo et al., 2020; Spagna et al., 2021, 2024; Pearson et al., 2015). Specific activations also emerged for each motivational state. For instance, bilateral activation of the cuneus was observed across all states except for “visceral” ones, whereas the left MTG was activated exclusively during “secondary” and “visceral” motivational states. The cuneus is implicated in imagination tasks but exhibits a negative correlation with the perceived vividness of mental imagery (Fulford et al., 2018). Together with the lingual gyrus, the cuneus is involved in both fundamental and higher-order visual processing functions, including orientation, direction, color perception, and face recognition. These regions also contribute to visual memory, creative thought, visual imagery, and language processing (Palejwala et al., 2021). As for the left MTG, it plays a crucial role in mental representation (Lima et al., 2016; Regev et al., 2021; Bartolomeo et al., 2020; Pearson et al., 2015), participating in social, emotional, and linguistic processing, as well as theory of mind—all of which are foundational to imaginative functions.

Affective states were associated with activations in the right fusiform gyrus and the left supramarginal gyrus. The fusiform gyrus plays a critical role in the perception and mental imagery of faces and bodies, as well as color processing. The importance of the right fusiform gyrus in face imagery is further underscored by clinical cases of patients with lesions in this region who experience prosopagnosia, impairing both perception and imagination of faces (Spagna et al., 2024). Additionally, affective states prompted activation in the supramarginal gyrus (BA40), situated at the temporoparietal junction. This area is part of the default mode network (DMN) and is implicated in self-awareness and self-referential thought (Doricchi, 2022), as well as in mental (Sasaoka et al., 2014) and spatial (Dijkstra et al., 2019) imagery. The recall of affective states also notably activated the right MFG, part of the anterior prefrontal region of interest (OBF). This same area was engaged in imagery tasks related to states of joy in the study by Proverbio et al. (2024). The OBF is integral to dopaminergic reward circuits (Floresco and Tse, 2007) and, by extension, to positive emotions (Proverbio et al., 2024). Specifically, Machado and Cantilino (2016) highlight that certain regions, including areas within the OBF, are uniquely active during experiences of happiness. Key regions associated with the sensation of joy include the mPFC, PCC, and the inferior parietal lobule. Moreover, “affective” states correspond to activations in the right superior temporal gyrus, which likely reflects the affective nature of stimuli within this category. This activation may also relate to the social nature of certain auditory stimuli that prompted imagery (Proverbio et al., 2024). Consistently, an fMRI study by Gawda et al. (2017) observed activation in frontal areas, including the right MFG, during a verbal fluency task involving “joyful words.” A distinct region associated with the “affective” motivational state was the right MTG, which, as also noted in Proverbio et al. (2024), is strongly linked to sadness. This area is activated in various contexts that evoke sadness, such as viewing sad faces or listening to sad music. Reduced activation in these regions has been observed in patients with impairments in recognizing affective information from faces or voices (Zuberer et al., 2022).

During the recall of “secondary” motivational states, heightened activation was observed in the right fusiform and MTG, regions associated with social play. The social component appears essential, as multiple studies implicate the temporal lobe in perceiving others’ behavior (Della Vedova and Proverbio, 2024). Additionally, the right superior temporal gyrus was activated during music imagery, a finding consistent with prior studies using visual imagery paradigms (Proverbio et al., 2023). This region, which includes Heschl’s gyrus and the auditory cortex, shows robust activation in tasks involving both music perception and imagination (Zatorre, 1999; Yu et al., 2017; Abraham, 2016; Zatorre et al., 2005; Regev et al., 2021). Studies on self-generated auditory imagery (Hu et al., 2023) demonstrate that both sound perception and its imaginative recreation activate primary and secondary auditory cortices. A further distinctive feature of “secondary” motivational states associated with movement involves the left precentral gyrus (Wei et al., 2022), playing a role in motor imagery (Ma et al., 2024), and the right MFG, which includes the supplementary motor area (SMA) (Yazawa et al., 2000) as well as parietal regions BA3 and BA7. Similarly, Della Vedova and Proverbio (2024) identified the premotor area, specifically in the left hemisphere, as crucial during motor imagery tasks. This area, in addition to being adjacent to the left precentral gyrus, functions in conjunction with it during movement execution (Urgesi et al., 2014).

During the recall of “somatosensory” states (such as cold or pain), the following areas emerged as distinctive: the right and left inferior parietal lobes, the right angular gyrus, and the right postcentral gyrus. The posterior parietal cortex, as well as the postcentral and intraparietal areas, appear to be particularly involved during both real and imagined tactile experiences (Chivukula et al., 2021). Moreover, the inferior parietal lobe is the lesion site in patients suffering from asymbolia for pain, highlighting its role as a key region for pain perception (Berlucchi and Vallar, 2018). The postcentral gyrus, corresponding to S1, receives tactile, nociceptive, thermal, and proprioceptive stimuli (Tamè et al., 2016), and therefore, strongly represents the imagined content recalled by participants during the study.

The recall of “visceral” motivational states (such as hunger and thirst) specifically activated the right MFG and left inferior frontal gyrus, linked to hunger sensations, as well as the right and left uncus, right precuneus, and right cingulate gyrus, involved in craving. Evidence supports the role of the frontal operculum (BA45) in gustatory processing (Veldhuizen et al., 2011), with Rolls et al. (1988) suggesting its involvement in continuous responses to gustatory stimuli rather than satiety. Notably, the right MFG is particularly active in patients with eating disorders during food-related tasks, supporting its role in food intake, though often with a pathological association (Celeghin et al., 2023). Conversely, Chen and Zeffiro (2020) identified the right MFG as more active in response to sweet food stimuli in healthy and overweight populations. The uncus and cingulate gyrus, part of the limbic system, are implicated in visceral and food-related cravings, with studies showing that stress-induced eating behaviors overstimulate limbic system neurons (Svetlak, 2010). The precuneus, active during craving states, shares similarities with food-seeking behavior, with Hartwell et al. (2011) showing its activation during nicotine craving in smokers. Functional connectivity in the DMN between the dorsal cingulate cortex and precuneus, along with retrosplenial cortex, is enhanced in anorexia nervosa patients, correlating with body image scores and suggesting a cognitive effort to control food desire, stronger than in healthy controls (Roger et al., 2022).

On the bases of the reviewed literature, it can therefore be argued that the areas identified as sources of the electrical activity captured by the EEG recording were consistent with, and reliably reflecting, the content of motivational states that participants were instructed to recall. Furthermore, it is confirmed that imagination induces neural engagement akin to that experienced during real, firsthand encounters. This study effectively identified specific ERP components associated with twelve motivational micro-categories—hunger, thirst, sleep, warmth, cold, pain, joy, sadness, fear, movement, music, and play— grouped within four broader macro-categories: visceral needs, somatosensory needs, affective states, and secondary needs. These findings hold valuable implications for advancing BCI research, particularly in assessing consciousness and uncommunicable needs in patients with disorders of consciousness. Key ERP markers, including the N400 (notably larger for affective and somatosensory states) and the P400 (more pronounced in visceral and secondary needs), offer insight into differentiating motivational states.

### Study limitations

However, as with many studies in this area, our research faces several limitations. First, the small sample size and restricted age range limit the generalizability of the findings; expanding the sample size and including more diverse age groups could enhance the robustness and applicability of these results. Additionally, only twelve motivational states were examined, though investigating a broader array of motivational states could yield important insights into the needs patients affected by consciousness disorder. Future research should aim to broaden the participant pool across a wider age range, including minors, adults aged 28–65, and elderly individuals (65+). Expanding the study to encompass other motivational states within each macro-category, such as emotional states like anger or disgust, and secondary needs like wanting to hear a story or be cared for, may be especially relevant for understanding everyday needs. Lastly, applying this framework to clinical populations, such as visually impaired individuals, may reveal meaningful differences in brain activity, further enriching the potential applications of this research. A further potential limitation might come from the fact that the imaginary motivational states were to be voluntary activated, and did not derive from real homeostatic needs (such as hunger or drug craving). This condition may not fully correspond to people’s experiences in real situations related to such needs, but the same criticality holds for any study involving imagery paradigms.

## Acknowledgements

We are grateful to Francesca Pischedda for her kind contribution to the experimental paradigm.

## Data-availability statement

All data generated or analyzed during this study are included in this published article and its supplementary information files. The data supporting the findings of this study are available from the corresponding author upon reasonable request. Due to privacy restrictions, some data may not be shared publicly.

